# Social and news media in mapping the endangered Mediterranean Monk seal: exceptional data gains but persistent bias

**DOI:** 10.1101/2025.01.20.633919

**Authors:** Andry Castro, César Capinha

**Affiliations:** Centro de Estudos Geográficos, Instituto de Geografia e Ordenamento do Território, Universidade de Lisboa, Lisboa, Portugal; Associated Laboratory Terra, Portugal

**Keywords:** Mediterranean monk seal, *Monachus monachus*, News media, Social media, Species distribution mapping

## Abstract

Social and news media are increasingly recognised as valuable sources of species distribution data, particularly for rare and endangered species. However, these data sources are prone to biases arising from uneven observation efforts, which pose challenges for their effective use. Despite this, the drivers and extent of such biases remain understudied. Here we investigate the contribution of social and news media to improving the availability of distribution data for a rare and endangered species, the Mediterranean Monk seal (*Monachus monachus*), in the Madeira and Porto Santo Islands. We also assess the relationship between the distribution data collected and socio-human factors likely to determine spatial and temporal variation in observation effort. We found that media sources provide a remarkable number of observation records for the species (n=302), which compared to the GBIF, iNaturalist and Observation.org combined (n=11), represent a 27.5-fold increase. The number of observations exhibited seasonal variation, with annual peaks occurring in the summer months. A generalised linear model assessing the spatial distribution of observation data also revealed a strong positive association with areas of high human concentration. Significant variables included population density, hotel density, and proximity to coastal recreational hotspots such as diving sites and bathing areas/non-open beaches. Our results underscore the potential of social and news media in documenting species distributions, but they also emphasise the importance of accounting for observational biases. Doing so is essential to ensure the reliable and effective application of social and news media sourced data in supporting species conservation efforts and management strategies.

## 1. INTRODUCTION

Unstructured data sources, such as online newspaper news, social media and blog posts, are becoming a relevant source of ecological and biological data (Jarić et al., 2020). This tendency is very much supported by advances in portability, autonomy and image acquisition capacity and quality in smartphones, widespread internet access and strong adoption of social media platforms (August et al., 2015). As of October 2024, over five billion of people were using social media (Datareportal, 2024). For instance, Facebook and Instagram have about 3 and 2 billion monthly active users, respectively, each potentially uploading photos and videos of relevance for ecological and conservation research. These massive sources of unstructured information are thus capable of increasing biodiversity knowledge (Jarić et al., 2020; Chowdhury et al., 2022; Chowdhury et al., 2023), a role of relevance in the current context of unprecedented loss of biodiversity (Dirzo et al., 2014; Ceballos et al., 2020; Cowie et al., 2022). Considering this enormous potential, these data can be key to support conservation efforts of endangered species (Toivonen et al., 2019), acting in pro-conservation human behaviour changes, increasing conservation funding, and new conservation-based legislation or governmental actions (Bergman et al., 2022).

A field of application where unstructured data sources, such as news and social media, have shown significant contributions in recent years is species distribution mapping (ElQadi et al., 2017). These platforms offer relevant opportunities to complement biogeographical knowledge by providing up to date, large-scale, and often cost-effective observation data. Previous studies have highlighted the potential of these sources to enhance knowledge about species’ areas of occurrence, distribution ranges, and range-shifting dynamics. One notable example is the use of social media to monitor the distribution of invasive species. Platforms like Facebook and Twitter have facilitated tracking the spread of lionfish (*Pterois miles*) in the Mediterranean Sea (Al Mabruk et al., 2020), or, for example, the Oak Processionary Moth (*Thaumetopoea processionea*) and Eastern Grey Squirrel (*Sciurus carolinensis*) (Daume, 2016). Also of potential relevance is the role that these sources can have in documenting the occurrence and distribution of rare, charismatic, and symbolic species, which often attract attention not only from specialists but also from the public. The sometimes striking appearance of these species, their ecological importance, or cultural significance increase their visibility in social media posts and news coverage, generating valuable datasets for conservation planning (Di Minin et al., 2015; Barve, 2014; Chowdhury et al., 2023; Chowdhury et al., 2024).

Despite their promise, data from these sources are not without challenges. Similar to findings in citizen science (Geldmann et al., 2016; Tiago et al., 2017), biases are likely to arise because social media and news coverage are closely tied to the spatial and temporal dynamics of human activity. For example, seasonal observation activity can lead to an overrepresentation of observations during peak periods, complicating efforts to assess species seasonal dynamics accurately (Rosário et al., 2024). Spatial biases further compound these challenges. Areas with continuous or seasonal concentrations of people, such as urban centres, tourist hotspots, and other accessible regions with features that attract human activity, are more likely to be overrepresented in opportunistic observational datasets (Tiago et al., 2017). This uneven spatial and temporal coverage by social and news media can distort the species patterns in collected data, potentially leading to erroneous assumptions about distribution, habitat preferences, or population density. To address these issues, a detailed understanding of the patterns and drivers of temporal and spatial observational biases, is needed.

Here, we investigate the contribution of social and news media to improving the availability of distribution data for a rare and endangered species, the Mediterranean Monk Seal (*Monachus monachus*), in the Madeira and Porto Santo Islands. The Mediterranean Monk Seal is an emblematic and charismatic species within the Madeira Autonomous Region and holds significant global conservation importance. It is classified as globally vulnerable (Karamanlidis et al., 2023). However, due to its extremely small population in the Madeira region, it should be considered critically endangered locally (Karamanlidis, 2024). In Madeira, the small subpopulation is primarily restricted to the Desertas Islands. Conservation efforts led by the regional government have been instrumental in recovering this population, which had declined to only 6–8 individuals in 1988 (Marchessaux, 1989). Recent estimates suggest a slow but steady recovery, with 21 individuals recorded in 2018 (Pires et al., 2020) and 27 in 2021 (Pires et al., 2023). As a result of these conservation successes, sightings of the Mediterranean Monk Seal have become more frequent on the main island of Madeira (Livramento, 2022). Understanding the spatial and temporal dynamics of the Mediterranean Monk seal in the region is thus essential for sustaining its recovery and guiding targeted conservation and management strategies. An Information Network has operated since 2002, gathering occasional sightings made from or reported to municipalities, whale-watching companies, and other entities (Pires et al., 2023). However, its reliance on opportunistic observation data highlights the need for complementary sources such as social and news media, which can broaden the spatial and temporal coverage and strengthen ongoing monitoring efforts.

We searched for and collected Mediterranean Monk Seal observations in the region by analysing publications from local news outlets and social media platforms, including *YouTube*, *Facebook*, *Instagram*, *blogs*, and *X*. We curated and integrated the collected information, considering the date of observation and its spatial precision. For reference, we also compared the volume and distribution of data collected with those from three popular biodiversity observation repositories: the Global Biodiversity Information Facility (GBIF; gbif.org), iNaturalist (inaturalist.org) and Observation.org (observation.org). Subsequently, we analysed the temporal dynamics and spatial distribution of the media-derived observations, evaluating whether the represented patterns could be attributed to human observation effort.

## 2. METHODOLOGY

**2.1 STUDY AREA**

The Madeira Autonomous Region is in the Atlantic Ocean, approximately 900 km southwest of mainland Portugal **(Fig. 1)**. It consists of two inhabited islands (Madeira and Porto Santo) and two uninhabited sub-archipelagos: the Selvagens Islands, situated 165 km north of the Canary Islands, and the Desertas Islands. The Desertas Islands, which include Bugio, Ilhéu Chão, and Deserta Grande, are the primary habitat of the Mediterranean Monk Seal. These islands are located 18 km southeast of Madeira Island, where the seals are observed with greater frequency.

**Figure 1.**
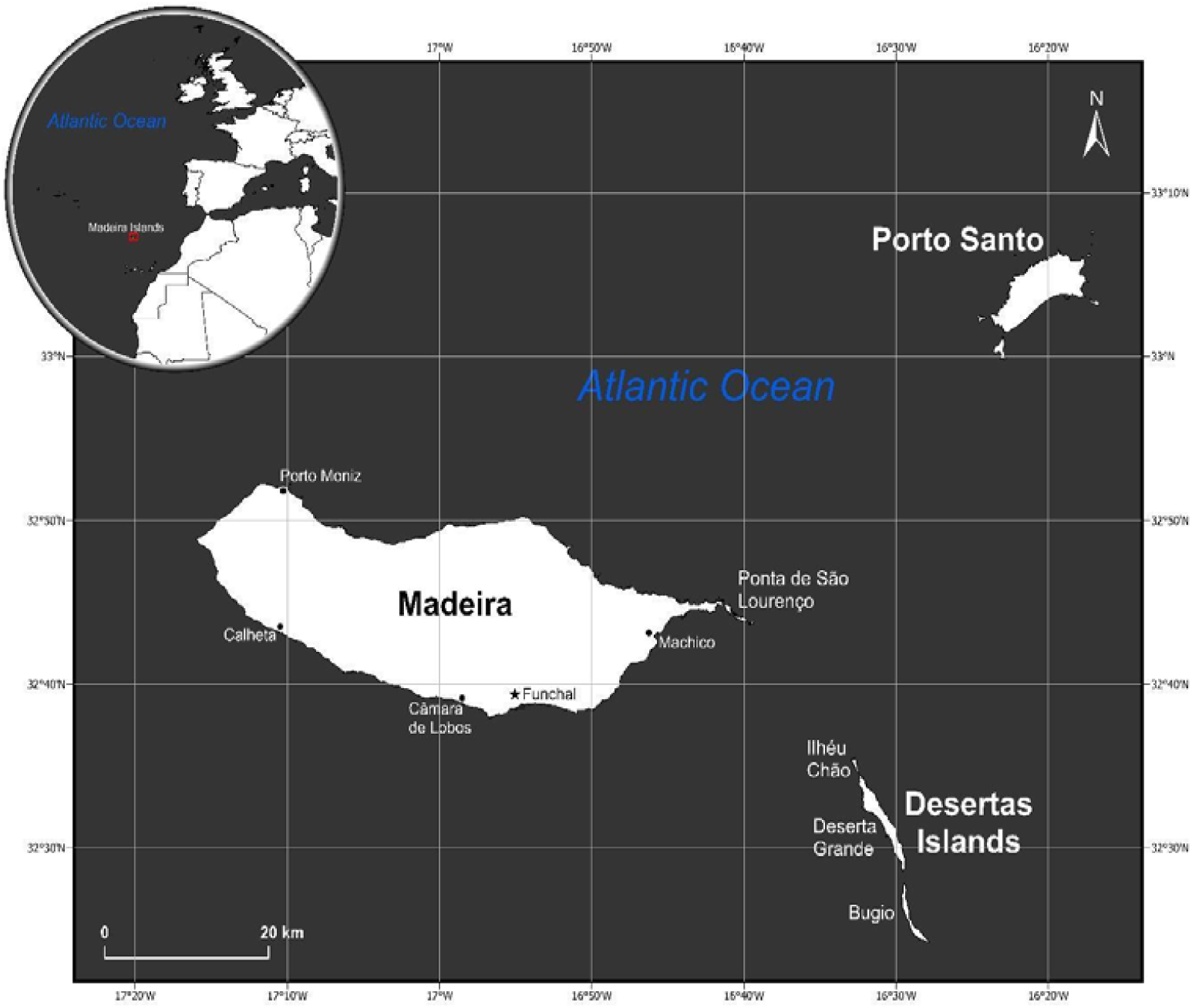
Map of the Madeira Autonomous Region. The inset shows the geographical position of the Madeira Islands in relation to European and African continents and the Atlantic Ocean. Towns and geographical features are labelled for reference.

### 2.2 OBSERVATION DATA COLLECTION

We searched for and collected data on spontaneous observations of the Mediterranean Monk Seal across the inhabited islands of the Madeira Autonomous Region (Madeira and Porto Santo) using media and social media platforms. The data collection focused on a 15-year temporal window, spanning from 2009 to November 2024.

From the news media, we considered the main news outlets of Madeira, which usually reports Monk seal sightings. The search was performed through extensive online browsing of newspaper archives focusing on the species common names in Portuguese (‘Lobo Marinho’). We only considered records that provided a minimum level of geographical detail for the sighting location, such as a region name, a specific spot, or an image with an identifiable location within the study area. Due to the replication dynamics of news media, we cleaned the collected data by deleting duplicates reporting the same sighting event and retaining the most detailed information across sources to obtain individual observation records of the species. We stored the title, description, URL, month, and year of the sighting reported in the news article.

For social media, we considered *YouTube, Facebook, Instagram, blogs*, and *X* within the same temporal window. Search in these platforms used both the species common names in English and in Portuguese (‘Lobo Marinho’; ‘Monk Seal’), in each social media platforms. As above, only records that provided identifiable geographical information relatable to the region were considered. For qualifying records, we stored the title, description (if available), URL, and the month and year of each record. To avoid duplicate reports of the same original sighting, we compared the distinct records from social media and cross-referenced them with those collected from the news media dataset. In cases of duplicate observations, we merged the records, prioritizing the most detailed information available from any of the sources and combining the data into a single record.

### 2.3 GEOREFERENCING OF OBSERVATION DATA

We georeferenced all collected observation records, providing decimal coordinates for the sighting location. Most social media posts and media reports allowed for a relatively precise identification of the observation location by reviewing the title and description of each post, and sometimes by analysing pictures, videos, and comments associated with the post. However, for some of the data, we could only assign approximate coordinates, as the social media and media information only provided coarse geographical information (e.g., the municipality name; e.g., ‘Funchal’) rather than a specific spot (e.g., Funchal harbour) and did not include images or videos for additional verification. Based on the spatial precision allowed by the geographical information provided, we classified records into one of three categories: spatial precision of 100m or better, between 100m and 500m, and less than 500m. Additionally, we added information on the corresponding parish and municipality names to all records.

### 2.4 BASELINE DATA FROM GBIF AND CITIZEN SCIENCE

To assess the data gains or complementarity of social media and news media in relation to conventional biodiversity observation data sources, we collected observation records of the species from GBIF and two well-known citizen science platforms, iNaturalist and Observation.org. For these three sources, we considered the same 15-year period (2009 to November 2024) and the same study area. For GBIF, we retrieved presence-only data with coordinates. For iNaturalist, we retrieved georeferenced observations with research-grade level (records supported by visual media, accurate location, date, and a community consensus on species identity). For Observation.org, we retrieved georeferenced records “accepted with evidence” (observations convincingly documented with images or sounds or approved by an appointed rarities committee), “accepted (by admin)” records (observations accepted based on expert knowledge, such as distribution, experience, or previous observations, but lacking documentation with images or sounds), and “accepted (automatic validation)” records (accepted through automated rules based on validated observations or image recognition).

### 2.5 DATA ON TEMPORAL AND SPATIAL PREDICTORS

A primary aim of our work was to examine whether the temporal and spatial patterns of observations collected from social and news media exhibited a signal of expected human observation effort. For that purpose, we collected several variables expected to represent patterns of spatial and temporal variation in this effort. For the temporal component, we collected data on tourist stays (‘guests number’) in tourist accommodation establishments by month). These data were sourced from Regional Directorate of Statistics of Madeira (DREM) and covered the period 2009 to 2023 with a monthly resolution. Our expectation is that this variable would capture the temporal dynamics of tourism presence on the islands, reflecting variability in social media coverage. Additionally, periods of higher tourist influx on the islands are also likely to correlate with a trend among residents to engage in outdoor and coastal activities, such as beach attendance and sport diving.

To explore potential drivers of spatial variation in observation effort, we collected a range of variables. These included human population density, hotels density, distance to ports, harbours and marinas, distance to the main diving spots used by sport diving companies, distance to open beaches (typically sand and pebble beaches), distance to balneary facilities and non-open beaches, and finally distance to the main streams mouths. A detailed characterisation of the spatial resolution, year of data and primary source for each is provided in **Table 1**.

**Table 1:** List of variables considered as predictors of spatial patterns of observation data collected from social and news media. (See Appendix A)

| Predictors | Spatial Resolution | Coordinate System | Base information |  |  |
| --- | --- | --- | --- | --- | --- |
|  |  |  | Description | Period | Source |
| Population density | Civil parish | PTRA08<br>UTM zone<br>28N | Resident population (No.) by locality | 2021 | National Institute for Statistics (INE) |
| Hotels density | 250 x 250 meters |  | Hotels in Madeira Autonomous Region | 2024 | Google maps extractor |
| Distance to Ports, harbours and marinas |  |  | Ports, harbours and marinas |  | World Imagery basemap from Environmental Systems Research Institute, Inc (ESRI) |
| Distance to balneary facilities and non-open beaches |  |  | Balneary facilities and non-open beaches |  |  |
| Distance to the main diving spots |  |  | Diving spots |  |  |
| Distance to the main streams mouths |  |  | Streams mouths |  |  |
| Distance to sand and pebble open beaches |  |  | Sand and pebble open beaches |  |  |

We hypothesised a positive and significant association between the probability of species recording and areas with higher human population density and hotel density. Conversely, we expected negative relationships with increasing distance to ports, harbours, marinas, balneary facilities, beaches, diving spots and main river mouths. Ultimately, these expectations reflect a straightforward hypothesis that the spatial patterns in collected observation records are primarily shaped by human observation effort.

### 2.6 STATISTICAL ANALYSIS

#### Temporal variation in number of observations

To assess the temporal variation in number of observations records we summed the number of records in each month of the year. We analysed both the cumulative number of records across years and their inter-annual monthly variation. To assess the degree of association between number of monthly observations and tourist stays, we calculated the Pearson and Spearman correlation coefficients using function ‘*rcorr’* from ‘*Hmisc’* package (Harrell, 2025).

#### Spatial distribution of observations

To analyse the spatial association between collected records and the spatial variables describing human observation effort, we focused on records having a spatial precision of 100 meters or better and located up to 50 meters inland and 200 meters out to sea (i.e., in the coastline). Applying these criteria allowed us to retain most records (n=172; 57%) while enhancing the reliability of our analysis by excluding spatially uncertain records and distant sea observations, where data on human observation effort (e.g., touristic vessel routes) is unavailable. Next, we aimed to represent the general conditions available within these areas to contrast with the observed patterns. To achieve this, we generated pseudo-absence records within the same 50-meter inland and 200-meter seaward buffer. To ensure a comprehensive representation of these conditions, we generated a number of pseudo-absence records five times greater than the number of observations (n=860).

We combined the two datasets, coding observations as 1 and pseudo-absences as 0. For each data point, we extracted the corresponding values of the predictor variables. To assess the relationships between the dependent variable and the predictors, we implemented a Generalised Linear Model (GLM). However, to avoid redundancy among predictors (defined as correlations of |0.7| or higher; Dormann et al., 2007), we first analysed their pairwise Pearson correlations using the ‘fuzzysim’ R package (Barbosa et al., 2024). Based on this analysis, the variable distance to ports, harbours, and marinas was excluded from the set of predictors. The GLM used a logit link function and a binomial error distribution to conform with expectations for binary response variables.

To assess the model’s ability to explain the spatial distribution of observation records as a function of supplied predictors, we performed a cross-validation procedure. Specifically, we partitioned the data into five folds, iteratively fitting the model with data from four partitions and comparing the predictions to the labels of the dependent variable in the remaining partition. This procedure was repeated five times, each time using a different partition for validation. To measure the agreement between predictions and validation data, we used the Boyce index (Boyce et al., 2002; Hirzel et al., 2006), which assesses how much model predictions differ from a random distribution of observed presences across prediction gradients. This metric is advantageous for evaluating presence-only data, as it does not consider pseudo-absences, which can be generated in locations of actual observations. A value of 1, or close to it, indicates that the location of observations predicted by the model closely align with the locations where they were observed. Conversely, values near zero suggest that the predicted spatial distribution resembles what would be expected by random chance. The GLM was built using R’s base ‘*glm’* function (R Core Team, 2023) and the Boyce index was calculated using the ‘*ecospat.boyce’* function from the ‘ecospat’ package (Broennimann et al., 2024).

## 3. RESULTS

### 3.1 DATA PATTERNS AND GAINS

A total of 302 Monk Seal observations were collected from social and news media. These data are distributed across all municipalities of Madeira Autonomous Region, 285 of which on the south coast of Madeira Island, 6 on the north coast and 9 on Porto Santo Island **(Fig. 2)**. The distribution across south coast is characterised by a large concentration of records along the urban axis Machico - Santa Cruz - Funchal - Câmara de Lobos (78% of all records), and the municipality of Calheta isolated in the southwest side of the Island (11.7%). Observations on Porto Santo Island are rare, totalling only 9 records. Most of the observations were collected from social media (n=254; 84.1%), while news media contributed a smaller but still relevant portion (n=48; 15.9%). Among social media platforms, Facebook was the primary source, accounting for 73.6% of the records, followed by Instagram (14.2%), YouTube (8.7%), blogs (2.4%), and *X* with 1.2%.

**Figure 2.**
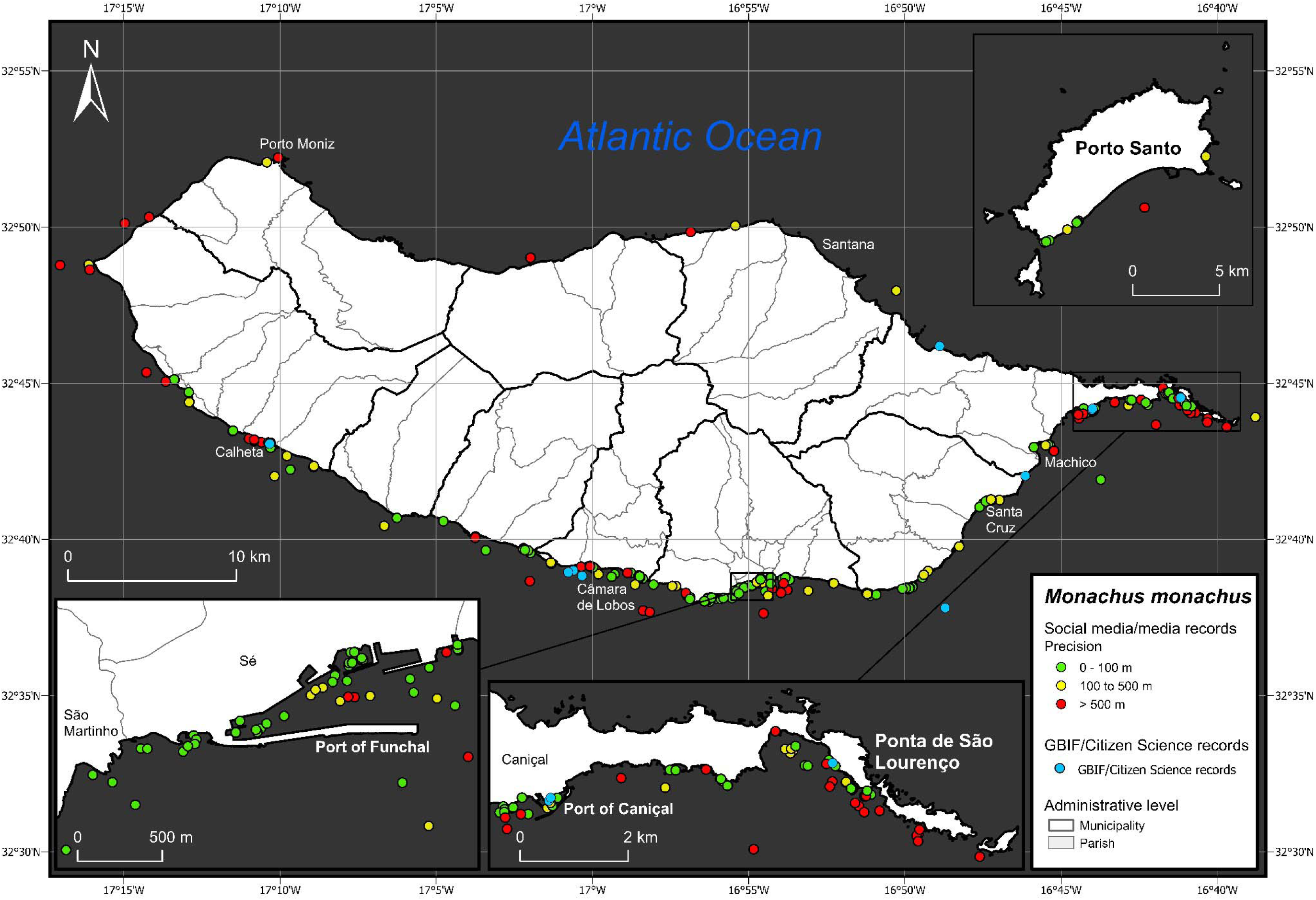
Distribution of Mediterranean Monk Seal observations in Madeira and Porto Santo Islands from 2009 to November 2024. The data are derived from publications in news outlets and social media, with additional observation records from GBIF and citizen science platforms (indicated in blue) included to evaluate data contributions. Observations from news and social media are color-coded from green to yellow to red, representing a gradient of decreasing estimated spatial precision. Administrative boundaries of municipalities and parishes are displayed to provide spatial context. (See **Appendix B**)

These numbers represent a substantial increase in data availability compared to the observations available in GBIF, iNaturalist, and Observation.org, which provided only 11 records for the same period. The spatial distribution of these database sourced records mirrors the general patterns represented by the social media and news-derived data, being predominantly concentrated along the south coast of Madeira Island, with a single record reported from the island’s northern region **(Fig. 2)**.

### 3.2 SEASONALITY OF RECORDINGS

The observations derived from news and social media reports show a marked seasonal trend, with records being substantially more frequent during the summer months **(Fig. 3)**. Specifically, 44.7% of all records occurred between July and September. In contrast, sightings are much less frequent during the autumn and winter months, with November and December having only 12 and 7 sightings, respectively, across the whole 15-year period. These patterns are moderately, albeit significantly correlated with the trends of visitation by tourists to the island (r=0.228; p = 0.002 and rL=0.280; p=0.0001).

**Figure 3.**
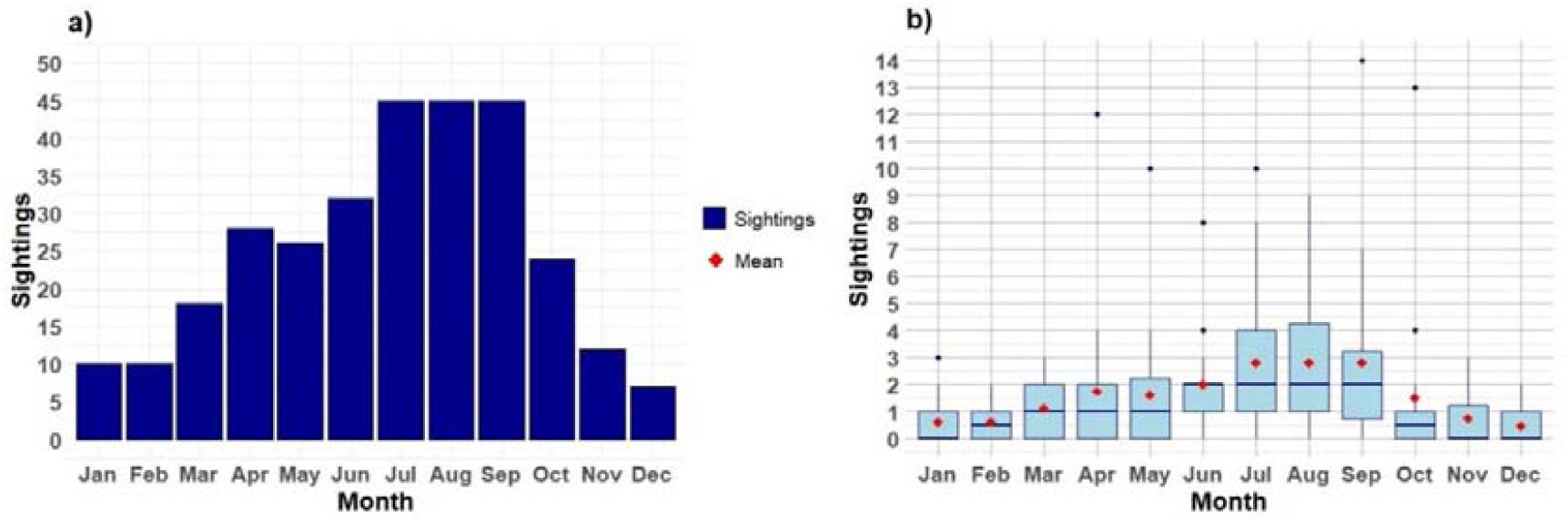
Temporal distribution of Mediterranean Monk seal observations collected from news and social media sources in Madeira and Porto Santo Islands between 2009 and November 2024.Panel (a) illustrates the total number of observations recorded for each month across all years. Panel (b) depicts the inter-annual distribution of observations by month.

### 3.3. PREDICTORS OF DISTRIBUTION

The GLM assessing the relationship between the spatial distribution of media-sourced observation records and proxies for human observation effort identified significant relationships for all predictors (p < 0.05), except for distance to beaches **(Table 2)**. The results align with expectations, showing a higher probability of observations in areas with greater human population density, higher hotel density, and proximity to balneary facilities/non-open beaches, major sport diving spots, and river stream mouths. Notably, the model demonstrated a good capacity to predict these patterns in left-out calibration data, achieving an average Boyce index value of 0.74 ± 0.12 across the five-fold cross-validation procedure.

**Table 2:** Results of the generalised linear model (GLM) explaining the spatial distribution of Mediterranean Monk Seal observation records, derived from social media and news sources, as a function of spatial descriptors associated with human activity and concentration propensity. Predictors with significant relationships (α = 0.05) have *p* values shown in bold.

| Predictor | Coefficient | Std. Error | $p$ value |
| --- | --- | --- | --- |
| Population density | 0.0003 | 0.0001113 | **0.005** |
| Hotels density | 0.5206 | 0.1054 | **<0.001** |
| Euclidean distance to balneary facilities and non-open beaches | -0.00002 | 0.00001116 | **0.042** |
| Euclidean distance to the main diving spots | -0.0004 | 0.00007202 | **<0.001** |
| Euclidean distance to the main streams mouths | -0.0002 | 0.00007112 | **0.027** |
| Euclidean distance to sand and pebble open beaches | -0.0001 | 0.00008491 | 0.474 |

## 4. DISCUSSION

We investigated the potential of social media and news outlets as sources of observation records for the endangered Mediterranean Monk Seal on Madeira and Porto Santo Islands. Our results revealed a substantial number of records available from these sources, greatly exceeding those found in established open-access biodiversity observation repositories such as GBIF, iNaturalist, and Observation.org. However, the collected data exhibited pronounced temporal and spatial patterns indicative of observational bias. Notably, these patterns showed significant correlations with various measures of human activity in the region, reflecting a putative influence of observation bias on available data.

Our results underscore that social and news media can serve as significant sources of distribution data, substantially surpassing popular biodiversity databases in coverage. In particular, the large degree of data gain compared to citizen-science platforms was somewhat unexpected, given that both involve opportunistic public contributions. This discrepancy indicates that citizen-science initiatives do not necessarily encompass the full potential of the general public to report biodiversity information. Instead, many social media users likely share observations of this charismatic species primarily for personal or recreational reasons, rather than with explicit scientific goals in mind, reflecting the Monk Seal’s charismatic appeal. In a broader context, this suggests that for species with similar public appeal, social and news media might outperform citizen-science platforms in terms of the quantity of observation data due to their widespread accessibility and more diverse user base.

The high number of records obtained indicate that data obtained solely from online media searches can serve as a valuable complement to formal monitoring programs. Several dedicated but non-public Monk Seal mapping efforts have also been conducted. For example, Pires et al. (2008) mapped 389 sightings reported to or collected by the Parque Natural da Madeira Service between 1988 and 2005. Additionally, an Information Network has been in operation since 2002, collecting Monk Seal sightings and imagery from occasional observers around Madeira Island (Pires et al., 2023). This network, which includes municipalities, whale-watching companies, and regional authorities, reported a total of 909 sightings between 2012 and 2021, though these presumably include duplicates. While the number of records we were able to collect from social and news media is lower, it is of the same order of magnitude and importantly captures the same overall spatial patterns as those identified in dedicated mapping efforts. For instance, as shown in Pires et al. (2008), our data similarly highlight the clustering of sightings in urban areas of Madeira Island, particularly between Machico and Câmara de Lobos, with noticeably fewer occurrences along the northern coast. These findings underscore the role of social and news media as a complementary approach for gathering distribution data on threatened and emblematic species. The role of these data may be even more important when sustained, formal mapping efforts are not feasible.

Importantly, however, observation records from social and news media must be evaluated for their representativeness before use. The patterns observed in opportunistic data result from the interaction of two main factors: first, the species’ true spatial dynamics, and second, extrinsic factors related to observation effort, which depend on when and where observers are present. Without precise measurements, it is nearly impossible to fully disentangle the relative influence of the latter factor on the patterns represented in the data. However, our approach -statistically relating temporal and spatial proxies of human effort to observation patterns -indicates that observational bias can be substantial.

Specifically, we identified significant associations between the spatial patterns of observations and multiple indicators of human activity concentration. Notably, a model relying solely on these indicators was able to predict the spatial distribution of observations with good accuracy. This suggests that the spatial distribution of observations is strongly influenced by unequal observation effort. While the precise movement dynamics of the species remain understudied, this conclusion is also supported by recent GPS collar data for two individuals (Pires et al., 2020). The data revealed extensive seal movement around Madeira Island, contrasting with the southernly clustered patterns represented in media-sourced data. As indicated by the results of our GLM, this pattern - at least to some extent - can likely be attributed to human demographic and economic factors. For example, the southern coast of Madeira is more densely populated and hosts significant economic activities and tourism, whereas the northern coast is only sparsely populated. These findings highlight the need for caution when using social and news media records for biogeographical purposes, such as assessing habitat usage or species distribution modelling.

We also detected a moderate but statistically significant relationship between monthly trends in the number of observations and tourist seasonality. It is plausible to expect greater social media coverage during periods of higher tourist activity, particularly in the summer months when larger numbers of people gather around the sea. This pattern of seasonality in observation numbers was indeed observed in our analysis. However, the extent to which this pattern also reflects the seasonality of Mediterranean Monk Seal occurrences is unclear. Interestingly, there are reasons to believe that this may partly represent true species activity. Annual activity in the Desertas Islands, the species’ primary breeding area, is highest during the autumn and winter months, driven by the timing of pupping events and post-parturition activities (Pires et al., 2007). Once pups are weaned, seals may change their behaviour, becoming more solitary and spending longer periods outside breeding areas, such as Madeira Island and Porto Santo, to meet dietary needs or rest in caves. To disentangle the specific contributions of observation effort versus true seasonal activity of the species in shaping the observed patterns, targeted research is required.

Despite the identified shortcomings, it is important to note that the potential biases identified are not unique to social and news media data. In our specific case, similar biases are likely to be present in structured datasets from citizen science platforms and even in dedicated mapping efforts that rely on opportunistic observation reporting. Relatedly, data sourced from social media and news outlets may jointly be used with data from these sources and also benefit from the application of analytical approaches commonly proposed to mitigate spatial biases in these sources (e.g., Fithian et al., 2015; Isaac et al., 2014).

Before considering the collation of observations from social and news media data, it is important to also recognize that this procedure can be time-consuming and resource-intensive. For example, search, filtering, and validation efforts were needed to remove irrelevant posts, such as those referencing the ferry “Lobo Marinho” or the Hawaiian Monk Seal (*Monachus schauinslandi*), both of which introduce “noise” to search results. Recent advancements in artificial intelligence, including large language models, can automate search result classification and information extraction (Castro et al., 2024). However, fully implementing these automated approaches requires specialised workflows that may be challenging to deploy across the multiple data sources. In addition, many social media posts lack precise geolocation data, leading to further effort in manual georeferencing. Multiple posts also often refer to the same event, creating duplicate records that need to be identified and merged. Ultimately, substantial effort can be required to produce a harmonized and robust dataset of available observations.

In conclusion, our findings demonstrate the potential of social and news media to complement traditional biodiversity monitoring efforts. However, biases resulting from non-random observation effort must be carefully considered before using the collected data. Advances in artificial intelligence offer promising tools to streamline data collation and processing, although challenges such as georeferencing are likely to persist and require considerable manual effort. Despite these limitations, social and news media data can be a highly relevant source of biogeographical data, particularly for species with strong public appeal. As the public continues to generate and share such data, its availability and utility in biodiversity mapping and monitoring are expected to grow even further over time.

## FUNDING

AC was supported by a grant (PRT/BD/152100/2021) financed by the Portuguese Foundation for Science and Technology, I.P. (FCT) under MIT Portugal Program. AC and CC acknowledge support from FCT through support to CEG/IGOT Research Unit (UIDB/00295/2020 and UIDP/00295/2020).

## ACKNOWLEDGMENTS

We would like to thank the constructive suggestions of the reviewers that helped to improve this work.

## SUPPLEMENTARY DATA

Appendix A – Predictors calculated: a) Distance to the main diving spots; b) Distance to balneary facilities and non-open beaches; c) Distance to Ports, harbours and marinas; d) Distance to the main streams mouths; e) Distance to sand and pebble open beaches; f) Hotels density; g) Population density by civil parish.

Appendix B – Mediterranean Monk Seal (*Monachus monachus*) records in Madeira and Porto Santo.

## CONFLICT OF INTEREST STATEMENT

No conflict of interest is declared.

## Notes

### Competing Interest Statement

The authors have declared no competing interest.

## REFERENCES

Al Mabruk, S. A. A., Rizgalla, J., Giovos, I., & Bariche, M. (2020). Social media reveals the first records of the invasive lionfish *Pterois miles* (Bennett, 1828) and parrotfish *Scarus ghobban* Forsskål, 1775 from Egypt (Mediterranean Sea). BioInvasions Records, 9(3), 574–579. 10.3391/bir.2020.9.3.13

August, T., Harvey, M., Lightfoot, P., Kilbey, D., Papadopoulos, T., & Jepson, P. (2015). Emerging technologies for biological recording. Biological Journal of the Linnean Society, 115(3), 731–749. 10.1111/bij.12534

Barbosa, A., Estrada, A., Melloy, P., & Guerrero, C. (2024). Fuzzy Similarity in Species Distributions. R package version 4.29. https://fuzzysim.r-forge.r-project.org/

Barve, V. (2014). Discovering and developing primary biodiversity data from social networking sites: A novel approach. Ecological Informatics, 24, 194–199. 10.1016/j.ecoinf.2014.08.008

Bergman, J., Buxton, R., Lin, H., Lenda, M., Attinello, K., Hajdasz, A., Rivest, S., Nguyen, T., Cooke, S., & Bennett, J. (2022). Evaluating the benefits and risks of social media for wildlife conservation. Facets, 7, 360–397. 10.1139/facets-2021-0112

Boyce, M. S., Vernier, P. R., Nielsen, S. E., & Schmiegelow, F. K. A. (2002). Evaluating resource selection functions. Ecological Modelling, 157, 281–300. 10.1016/S0304-3800(02)00200-4

Broennimann, O., Di Cola, V., Petitpierre, B., Breiner, F., Scherrer, D., D’Amen, M., Randin, C., Engler, R., Hordijk, W., Mod, H., Pottier, J., Di Febbraro, M., Pellissier, L., Pio, D., Garcia Mateo, R., Dubuis, A., Maiorano, L., Psomas, A., Ndiribe, C., … & Guisan, A. (2024). Spatial ecology miscellaneous methods. R package version 4.1.1. https://www.unil.ch/ecospat/home/menuguid/ecospat-resources/tools.html

Castro, A., Pinto, J., Reino, L., Pipek, P., & Capinha, C. (2024). Large language models overcome the challenges of unstructured text data in ecology. Ecological informatics, 82, 102742. 10.1016/j.ecoinf.2024.102742

Ceballos, G., Ehrlich, P. R., & Raven, P. H. (2020). Vertebrates on the brink as indicators of biological annihilation and the sixth mass extinction. Proceedings of the National Academy of Sciences, 117(24), 13596–13602. 10.1073/pnas.1922686117

Chowdhury, S., Aich, U., Rokonuzzaman, M., Alam, S., Das, P., Siddika, A., … & Callaghan, C. T. (2023). Increasing biodiversity knowledge through social media: A case study from tropical Bangladesh. BioScience, 73(6), 453–459. 10.1093/biosci/biad042

Chowdhury, S., Aich, U., Rokonuzzaman, M., Alam, S., Das, P., Siddika, A., … & Fuller, R. A. (2022). *Spatial occurrence data for the animals of Bangladesh derived from Facebook* (Dataset). PANGAEA. https://doi.pangaea.de/10.1594/PANGAEA.948104

Chowdhury, S., Hawladar, N., Roy, R. C., Capinha, C., Cassey, P., Correia, R. A., Deme, G. G., Di Marco, M., Di Minin, E., Jarić, I., Ladle, R. J., Lenoir, J., Momeny, M., Rinne, J. J., Roll, U., & Bonn, A. (2024). Harnessing social media data to track species range shifts. EcoEvoRxiv. 10.32942/X2R63N

Cowie, R. H., Bouchet, P., & Fontaine, B. (2022). The Sixth Mass Extinction: Fact, fiction or speculation? Biological Reviews, 97(2), 640–663. 10.1111/brv.12816

Daume, S. (2016). Mining Twitter to monitor invasive alien species — An analytical framework and sample information topologies. Ecological Informatics, 31, 70–82. 10.1016/j.ecoinf.2015.11.014

DataReportal (2024, January 10). Global social media statistics. DataReportal. https://datareportal.com/social-media-users

Di Minin, E., Tenkanen, H., & Toivonen, T. (2015). Prospects and challenges for social media data in conservation science. Frontiers in Environmental Science, 3, Article 63. 10.3389/fenvs.2015.00063

Dirzo, R., Young, H., Galetti, M., Ceballos, G., Isaac, N., & Collen, B. (2014). Defaunation in the Anthropocene. Science, 345(6195), 401–406. 10.1126/science.1251817

Dormann, C. F., McPherson, J. M., Araújo, M. B., Bivand, R., Bolliger, J., Carl, G., Davies, R. G., Hirzel, A., Jetz, W., Kissling, W. D., Kühn, I., Ohlemüller, R., Peres-Neto, P. R., Reineking, B., Schröder, B., Schurr, F. M., & Wilson, R. (2007). Methods to account for spatial autocorrelation in the analysis of species distributional data: A review. Ecography, 30(5), 609–628. 10.1111/j.2007.0906-7590.05171.x

ElQadi, M. M., Dorin, A., Dyer, A., Burd, M., Bukovac, Z., & Shrestha, M. (2017). Mapping species distributions with social media geo-tagged images: Case studies of bees and flowering plants in Australia. Ecological Informatics, 39, 23–31. 10.1016/j.ecoinf.2017.02.006

Fithian, W., Elith, J., Hastie, T., & Keith, D. A. (2015). Bias correction in species distribution models: Pooling survey and collection data for multiple species. Methods in Ecology and Evolution, 6(4), 424–438. 10.1111/2041-210X.12242

Geldmann, J., Heilmann-Clausen, J., Holm, T., Levinsky, I., Markussen, B., Olsen, K., Rahbek, C., & Tøttrup, A. (2016). What determines spatial bias in citizen science? Exploring four recording schemes with different proficiency requirements. Diversity and Distributions, 22(11), 1139–1149. 10.1111/ddi.12477

Harrell, F. E. Jr., & Dupont, C. (2024). Harrell Miscellaneous. R package version 5.2-2. https://hbiostat.org/r/hmisc/

Hirzel, A. H., Le Lay, G., Helfer, V., Randin, C., & Guisan, A. (2006). Evaluating the ability of habitat suitability models to predict species presences. Ecological Modelling, 199(2), 142–152. 10.1016/j.ecolmodel.2006.05.017

Isaac, N.J.B., van Strien, A.J., August, T.A., de Zeeuw, M.P. & Roy, D.B. (2014) Statistics for citizen science: extracting signals of change from noisy ecological data. Methods in Ecology and Evolution, 5, 1052–1060.

Jarić, I., Correia, R. A., Brook, B. W., Buettel, J. C., Courchamp, F., di Minin, E., Firth, J. A., Gaston, K. J., Jepson, P., Kalinkat, G., Ladle, R., Soriano-Redondo, A., Souza, A. T., & Roll, U. (2020). iEcology: Harnessing large online resources to generate ecological in sights. Trends in Ecology and Evolution, 35(7), 630–639. 10.1016/j.tree.2020.03.003

Karamanlidis, A. (2024). Current status, biology, threats and conservation priorities of the Vulnerable Mediterranean monk seal. Endangered Species Research, 53, 341–361. 10.3354/esr01304

Karamanlidis, A.A., Dendrinos, P., Fernandez de Larrinoa, P., Kıraç, C.O., Nicolaou, H. & Pires, R. (2023). Monachus monachus. The IUCN Red List of Threatened Species 2023: e.T13653A238637039. 10.2305/IUCN.UK.2023-1.RLTS.T13653A238637039.en (accessed 11 Dec 2023)

Livramento, M. (2022, 17 de agosto). Lobo-marinho avistado mais vezes na Madeira. Diário de Notícias Madeira. https://www.dnoticias.pt/2022/8/17/324372-lobo-marinho-avistado-mais-vezes-na-madeira/

Marchessaux, D. (1989). Distribution et statut des populations du phoque moine Monachus monachus (Hermann, 1779). Mammalia, 53(4), 621–628. 10.1515/mamm.1989.53.4.62114

Pires, R., Costa Neves, H., & Karamanlidis, A. (2007). Activity Patterns of the Mediterranean Monk Seal (Monachus monachus) in the Archipelago of Madeira. Aquatic Mammals, 33(3),327–336. 10.1578/AM.33.3.2007.327

Pires, R., Costa Neves, H., & Karamanlidis, A. (2008). The Critically Endangered Mediterranean monk seal Monachus monachus in the archipelago of Madeira: priorities for conservation. Oryx, 42(2), 278–285. 10.1017/S0030605308006704

Pires, R., Aparicio, F., & Fernández de Larrinoa, P. (2020). Estratégia para a conservação do lobo-marinho no Arquipélago da Madeira. Instituto das Florestas e Conservação da Natureza. https://ifcn.madeira.gov.pt/images/Doc_Artigos/Biodiversidade/Projetos/lobomarinho/Estrategia_para_a_Conservacao_do_Lobo-marinho_na_Madeira.pdf

Pires, R., Aparicio, F., Baker, J. D., Pereira, S., Caires, N., Cedenilla, M. A., Harting, A. L., Menezes, D., & Fernández de Larrinoa, P. (2023). First demographic parameter estimates for the Mediterranean monk seal population at Madeira, Portugal. Endangered Species Research, 51, 269–283. 10.3354/esr01260

R Core Team (2023). R: A language and environment for statistical computing. R Foundation for Statistical Computing, Vienna, Austria. URL: https://www.R-project.org/.

Rosário, I. T., Tiago, P., Chozas, S., & Capinha, C. (2024). Drivers of temporal bias in biodiversity recording by citizen scientists. EcoEvoRxiv. 10.1101/2024.07.22.604598

Tiago, P., Ceia-Hasse, A., Marques, T. A., Capinha, C., & Pereira, H. M. (2017). Spatial distribution of citizen science casuistic observations for different taxonomic groups. Scientific reports, 7(1), 12832. 10.1038/s41598-017-13130-8

Toivonen, T., Heikinheimo, V., Fink, C., Hausmann, A., Hiippala, T., Järv, O., Tenkanen, H., & Di Minin, E. (2019). Social media data for conservation science: A methodological overview. Biological Conservation, 233, 298–315. 10.1016/j.biocon.2019.01.023

