## Supplementary figures and images for "Social and news media in mapping the endangered Mediterranean Monk seal: exceptional data gains but persistent bias"

### Appendix A

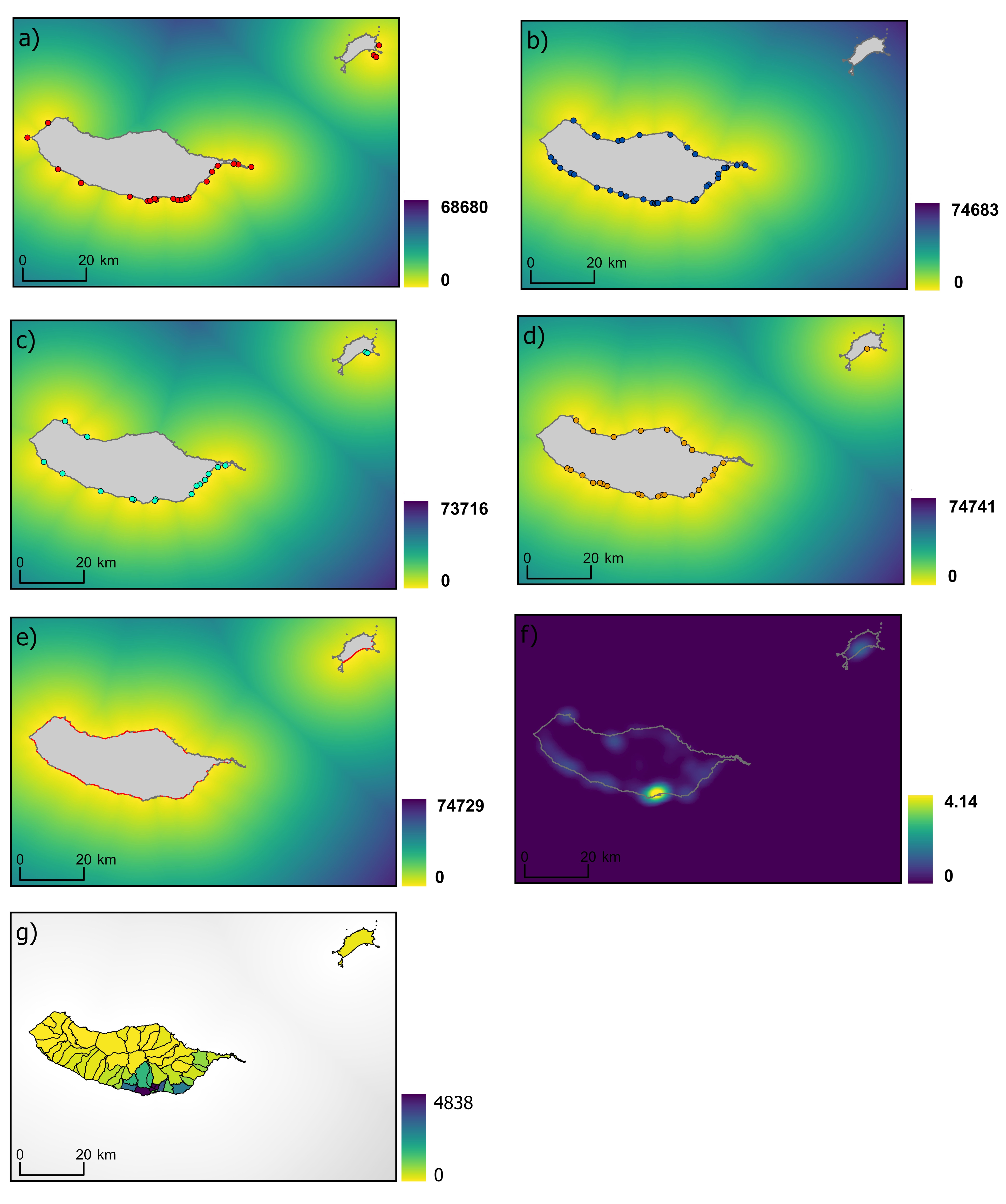
